# Oo-site: A dashboard to visualize gene expression during *Drosophila* oogenesis reveals meiotic entry is regulated post-transcriptionally

**DOI:** 10.1101/2022.01.31.478569

**Authors:** Elliot T. Martin, Kahini Sarkar, Alicia McCarthy, Prashanth Rangan

## Abstract

Determining how stem cell differentiation is controlled has important implications for understanding the etiology of degenerative disease and designing regenerative therapies. *In vivo* analyses of stem cell model systems have revealed regulatory paradigms for stem cell self-renewal and differentiation. The germarium of the female *Drosophila* gonad, which houses both germline and somatic stem cells, is one such model system. Bulk mRNA sequencing (RNA-seq), single-cell (sc) RNA-seq, and bulk translation efficiency of mRNAs are available for stem cells and their differentiating progeny within the *Drosophila* germarium. However, visualizing those data is hampered by the lack of a tool to spatially map gene expression and translational data in the germarium. Here, we have developed Oo-site (https://www.ranganlab.com/Oo-site), a tool for visualizing bulk RNA-seq, scRNA-seq, and translational efficiency data during different stages of germline differentiation, that makes these data accessible to non-bioinformaticians. Using this tool, we recapitulated previously reported expression patterns of developmentally regulated genes and discovered that meiotic genes, such as those that regulate the synaptonemal complex, are regulated at the level of translation.

## Introduction

The *Drosophila* ovary provides a powerful system to study stem cell differentiation *in vivo* (Bastock and St Johnston, 2008; Eliazer and Buszczak, 2011; Lehmann, 2012; Spradling et al., 2011). The *Drosophila* ovary consists of two main cell lineages, the germline, which ultimately gives rise to eggs, and the soma, which surrounds the germline and plays a supportive role in egg development (Eliazer and Buszczak, 2011; Roth, 2001; Schüpbach, 1987; Xie and Spradling, 2000). Each stage of *Drosophila* female germline stem cell (GSC) differentiation is observable and identifiable, allowing GSC development to be easily studied (Bastock and St Johnston, 2008; Lehmann, 2012; Xie and Spradling, 1998). Specifically, female *Drosophila* GSCs undergo an asymmetric division, giving rise to another GSC and a cystoblast (CB) (**Figure 1A**) (Chen and McKearin, 2003b; McKearin and Ohlstein, 1995; Xie and Spradling, 1998). The GSC and CB are marked by a round structure called the spectrosome (**Figure 1A**) (De Cuevas and Spradling, 1998; Zaccai and Lipshitz, 1996). The CB then undergoes four incomplete divisions resulting in 2-, 4-, 8-, and finally 16-cell cysts (CC), which are marked by an extended structure called the fusome (**Figure 1A**) (Chen and McKearin, 2003a, 2003b; De Cuevas and Spradling, 1998). In the 16-CC, one of the cyst cells is specified as the oocyte, while the other 15 cells become nurse cells, which provide proteins and mRNAs to support the development of the oocyte (**Figure 1A**) (Bastock and St Johnston, 2008; Carpenter, 1975; Huynh and St Johnston, 2000, 2004; Navarro et al., 2001; Theurkauf et al., 1993). The 16-CC is encapsulated by somatic cells and buds off from the germarium, forming an egg chamber (**Figure 1A**) (Bastock and St Johnston, 2008; Forbes et al., 1996; Xie and Spradling, 2000). In each chamber, the oocyte grows as the nurse cells synthesize and then deposit mRNAs and proteins into the oocyte, which eventually gives rise to a mature egg (Bastock and St Johnston, 2008; Huynh and St Johnston, 2000).

**Figure 1:**
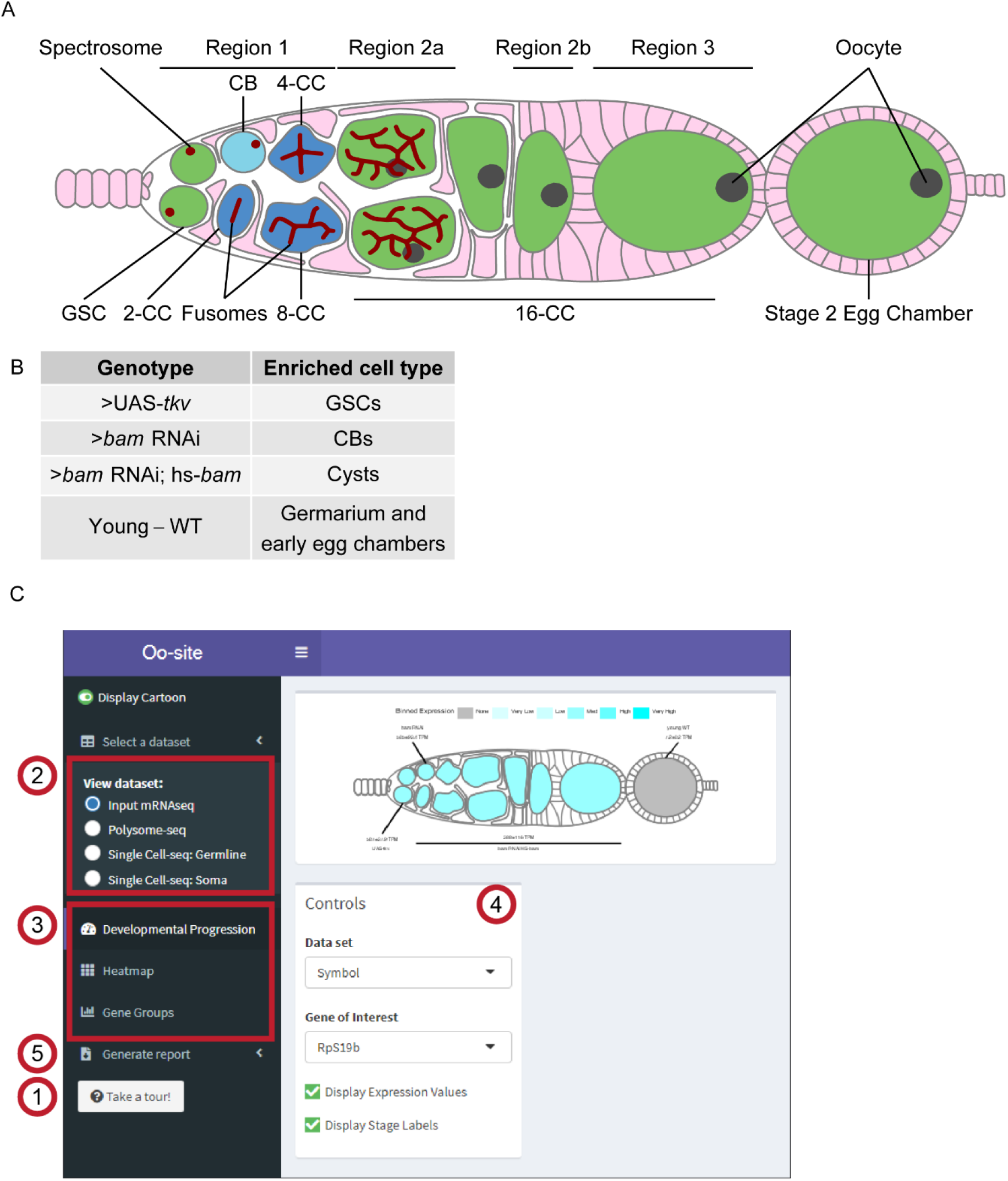
Oo-site integrates and provides an interface for interacting with multi-omic data covering major stages of *Drosophila* GSC differentiation. (**A**) Schematic illustrating developmental stages of germline development. (**B**) Summary of the samples used for bulk RNA-seq and polysome-seq and the cell types these samples are enriched for. (**C**) Screenshot of Oo-site dashboard, indicating: (1) “Take a Tour!” function, which guides the user through the functionality and operation of Oo-site. (2) The available seq datasets which the user can view, including RNA-seq of ovaries genetically enriched for developmental stages (bulk RNA-seq), polysome-seq of ovaries genetically enriched for developmental stages (Polysome-seq), single-cell seq of germline stages (Single-Cell seq: Germline), and single-cell seq of somatic stages in the germarium (Single-Cell seq: Soma). (3) the available visualizations which the user can use, including viewing the expression of genes over development at the level of a single gene (Developmental Progression), viewing all significantly changing genes as heatmaps (Heatmap), and viewing groups of genes either derived from GO-term categories or supplied by the user (Gene Groups). (4) The control panel, which the user can use to control the current visualization, and (5) the Generate Report Function, which can be used to download a PDF report of either the current visualization or all active visualizations.

Expression of differentiation factors, including those that regulate translation, results in progressive differentiation of GSCs to an oocyte (Blatt et al., 2020; Slaidina and Lehmann, 2014). In the CB, Bag-of-marbles (Bam) expression promotes differentiation and the transition from CB to 8-CC stage (Chen and McKearin, 2003a; McKearin and Ohlstein, 1995; Ohlstein and McKearin, 1997). In the 8-CC, RNA-binding Fox protein 1 (Rbfox1) promotes exit from the mitotic cell cycle into meiosis (Carreira-Rosario et al., 2016). Both the differentiation factors Bam and Rbfox1 affect the translation of mRNAs to promote differentiation (Carreira-Rosario et al., 2016; Li et al., 2009; Tastan et al., 2010). In addition, in 8-CCs, recombination is initiated across many cyst cells and then eventually is restricted to the specified oocyte (Hinnant et al., 2020; Huynh and St Johnston, 2000). Neither the mRNAs that are translationally regulated during this progressive differentiation nor how recombination is temporally regulated is fully understood (Cahoon and Hawley, 2016; Carreira-Rosario et al., 2016; Flora et al., 2018; Rubin et al., 2020; Slaidina and Lehmann, 2014; Tanneti et al., 2011; Wei et al., 2014).

Within the germarium, the germline is surrounded by and relies on distinct populations of somatic cells for signaling, structure, and organization (Roth, 2001; Schüpbach, 1987; Xie and Spradling, 2000, 1998). For example, the terminal filament, cap, and anterior-escort cells act as a somatic niche for the GSCs (Decotto and Spradling, 2005; Lin and Spradling, 1993; Wang and Page-McCaw, 2018; Xie and Spradling, 2000). Once GSCs divide to give rise to CBs, posterior escort cells guide CB differentiation by encapsulating the CB and the early-cyst stages (Kirilly et al., 2011; Shi et al., 2021; Upadhyay et al., 2016). Follicle stem cells (FSCs), which are present towards the posterior of the germarium, divide and differentiate to give rise to follicle cells, (FCs) which surround the late-stage cysts that give rise to egg chambers (Margolis and Spradling, 1995; Nystul and Spradling, 2010; Rust et al., 2020). FSCs also give rise to stalk cells and polar cells which connect the individual egg chambers that comprise the ovariole (Margolis and Spradling, 1995; Nystul and Spradling, 2010; Rust et al., 2020; Sahai-Hernandez et al., 2012).

While there is a wealth of bulk RNA-seq, single-cell mRNA-seq (scRNA-seq), and translational efficiency data from polysome-seq experiments for the cells in the germarium, there are several hurdles for easy utilization of this data:

1. scRNA-seq has exquisite temporal resolution but it can miss some lowly expressed transcripts which are better captured by bulk RNA-seq (Lähnemann et al., 2020). However, there is no easy way to compare these two data sets.
2. While scRNA-seq provides mRNA levels, it does not indicate if these mRNAs are translated, especially in the germline where translation control plays an important role (Blatt et al., 2020; Slaidina and Lehmann, 2014).
3. Lastly, there is a barrier to the visualization of the data for those who are not experienced in bioinformatics.

Here, we have developed a tool that we call Oo-site which integrates bulk RNA-seq, scRNA-seq, and polysome-seq data to spatially visualize gene expression and translational efficiency in the germarium.

## Results

To make bulk RNA-, scRNA-, and polysome-, seq data accessible to the community, we have collated and reprocessed previously published sequencing datasets of ovaries enriched for GSCs, CBs, cysts, and egg chambers (**Figure 1B**). Notably, each genetically enriched sample had matched bulk RNA-seq and polysome-seq libraries prepared, allowing for simultaneous read-out of mRNA level and translation status (**Supplemental Figure 1A**). One limitation is that the enriched cyst stages do not resolve each distinct stage of cyst development, instead, these samples represent a mixture of cyst stages. Therefore to supplement the enrichment data, we have integrated scRNA-seq data from Slaidina *et al*. which provides a more discrete temporal resolution of the cyst stages (Slaidina et al., 2021). We present these data as a tool called Oo-site (https://www.ranganlab.com/Oo-site), a collection of interactive visualizations that allows researchers to easily input a gene or collection of genes of interest to determine their expression pattern(s).

Oo-site consists of three modules: ovary-map, ovary-heatmap, and ovary-violin (**Figure 1C**). Each module allows users to visualize expression from matched mRNA-seq and polysome-seq data of genetically enriched stages of early GSC differentiation as well as previously published scRNA-seq data (Slaidina et al., 2021). Additionally, we have integrated scRNA-seq expression data for genes that cluster in somatic cell populations that reside in the germarium (Slaidina et al., 2021), however, here we focus on the germline (Slaidina et al., 2021). Ovary-map allows users to visualize the expression of a single gene over the course of differentiation in the framework of a germarium schematic, which contextualizes staging for those less familiar with *Drosophila* oogenesis. Ovary-heatmap consists of a clustered, interactive heatmap of genes determined to be differentially expressed that allows users to explore expression trends over-development (**Figure 1B, Supplemental Figure 1B-C’**). Finally, ovary-violin allows users to visualize the expression of multiple genes over the course of differentiation (**Figure 1C**). These groups of genes can be selected either by a GO-term of interest or a custom list of genes supplied by the user. The user can download a spreadsheet of gene expressions corresponding to the subset of selected or input genes. Finally, Oo-site incorporates a reporting tool that generates a downloadable report of the visualization(s) in a standardized format to facilitate their use for publication (**Figure 1C**). Researchers can use these datasets to enhance hypothesis generation or to confirm expression patterns observed from other methods.

Using Oo-site, we first determined if the bulk RNA-seq data that was acquired by enriching for specific stages of germline development is representative of the gene expression patterns from purified cell types. We compared bulk RNA-seq data obtained by enriching for GSC and CB cell types without purification from somatic cells (**Figure 1C**) to the GSC and CB data from Wilcockson *et al*. where they included a fluorescent-assisted cell sorting (FACS) step to eliminate somatic cells so that a pure population of these germline cells was sequenced (Wilcockson and Ashe, 2019). We analyzed the expression of genes that Wilcockson *et al*. identified as 2-fold or more down-or upregulated with a p-value < 0.01. We found that in the enriched bulk RNA-seq data these genes followed similar trends as identified by Wilcockson *et al*., indicating that despite the lack of FACS purification, enrichment of cell types reproduces meaningful mRNA expression changes over these stages (**Supplemental Figure 2A-A’**).

To determine if the bulk RNA-seq data recapitulates genuine changes in gene expression, we compared the expression of *ribosomal small subunit protein 19b* (*RpS19b*) in bulk RNA-seq to scRNA-seq data. Our bulk RNA-seq data, as well as the available scRNA-seq data indicated that *RpS19b* was highly expressed in GSCs, decreased during differentiation in the cyst stages and was greatly decreased in expression in early egg chambers, consistent with previous reports (**Fig 2A-B**) (McCarthy et al., 2021; Sarkar et al., 2021). To further validate this expression pattern, we probed the expression of *RpS19b in vivo* using *in situ* hybridization as well as an RpS19b::GFP line that is under endogenous control elements (McCarthy et al., 2021). We found that *RpS19b* was present in the GSCs and diminishes in the cyst stages both at the mRNA and protein level (**Figure 2C-E’**). Additionally, RpS19b::GFP expression resembled its mRNA expression indicating that its dynamic expression is achieved primarily through modulating the mRNA level of *RpS19b*, consistent with its moderate to high translational efficiency in early stages (**Figure 2C-D, Supplemental Figure 2B**). Thus, enriching for specific germline stages captures changes to gene expression in the germline. However, we note that care should be taken in interpreting bulk RNA-seq results as the data may be influenced by the somatic cells present in the samples. However, simultaneous comparison with scRNA-seq can alleviate this problem.

**Figure 2:**
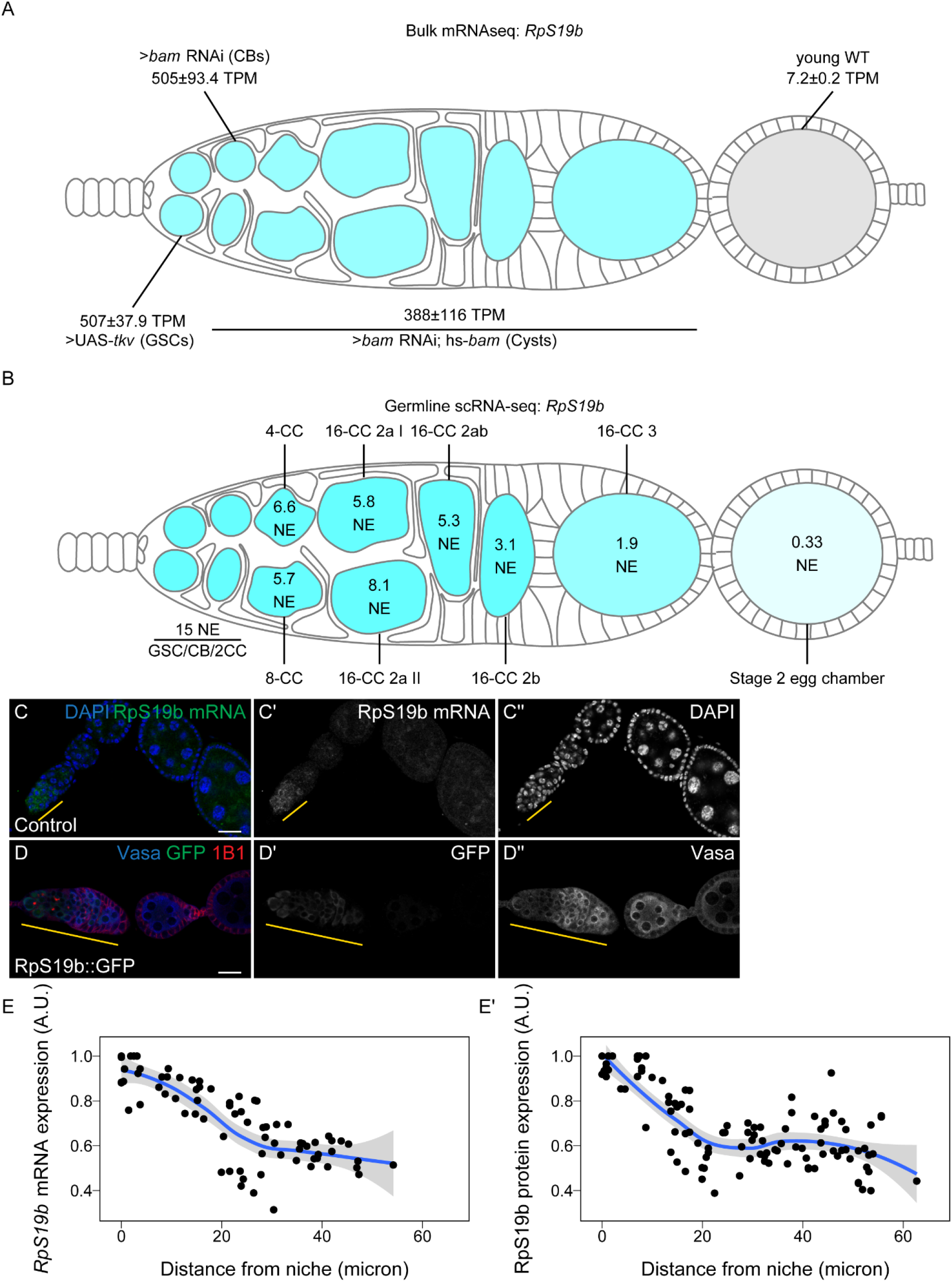
Oo-site allows for visualization of dynamically regulated genes. (**A-B**) Visualization of expression of *RpS19b* over germline development from (A) developmentally enriched stages and (B) single-cell seq data indicate that the mRNA level of *RpS19b* decreases starting in the cysts and is dramatically decreased in early egg chambers. Color indicates relative expression and values indicate the (A) mean TPM±standard error or (B) the normalized expression of *RpS19b* in each given stage. (**C-C’’**) Confocal images of ovaries with *in situ* hybridization of *RpS19b* (green, middle greyscale) and stained for DAPI (blue, right greyscale) demonstrate that the mRNA level of *RpS19b* decreases starting in the cyst stages and are dramatically lower in early egg chambers consistent with the seq data. (**D-D’’**) Confocal images of ovaries expressing RpS19b::GFP, visualizing (D’) GFP (green, middle greyscale), (D’’) Vasa staining (blue, right greyscale), and 1B1 (red) demonstrate that the protein expression of RpS19b::GFP is consistent with its mRNA levels. (**E-E’**) Quantifications of normalized mean intensity of staining, X-axis represents the distance in microns from the niche, Y-axis represents mean intensity normalized to the maximum mean intensity per germarium of (E) *RpS19b* mRNA or (E’) RpS19b::GFP. The line represents fit using a loess regression, shaded area represents the standard error of the fit. (n=5 germaria).

To determine the groups of genes that change as the GSCs differentiate into an egg, we used gene ontology (GO)-term analysis to probe for pathways that change at the level of RNA using bulk RNA-seq data. We did not identify any significant GO-terms in genes that are differentially expressed between GSCs and CBs. We found that genes with lower expression in GSCs compared to differentiating cysts are enriched in the GO-term polytene chromosome puffing which is consistent with GO-terms identified in Wilcockson *et al*. for genes that are expressed at lower levels in GSCs than in differentiating cysts than GSCs (**Figure 3A**). We also identified the polytene chromosome puffing GO-term in genes downregulated in CBs compared to cysts. Additionally, we observed that several GO-terms involving peptidase activity were enriched in genes upregulated in GSCs and CBs compared to cysts (**Figure 3B**). This is consistent with findings suggesting that peptidases can be actively regulated during differentiation and can influence stem cell fate (Han et al., 2015; Perišić Nanut et al., 2021; Tiaden et al., 2012). We found that two GO-terms related to glutathione transferase activity were enriched in genes downregulated in GSCs and CBs compared to ovaries from young-wildtype (young-WT) flies and in CBs compared to differentiating cysts, suggesting that metabolic processes may be altered during GSC differentiation. Additionally, comparison of CBs and differentiating cysts to young-WT, which contain early egg chambers, indicated that downregulated genes were enriched in GO-terms involving vitelline and eggshell coat proteins (**Figure 3A**).

**Figure 3:**
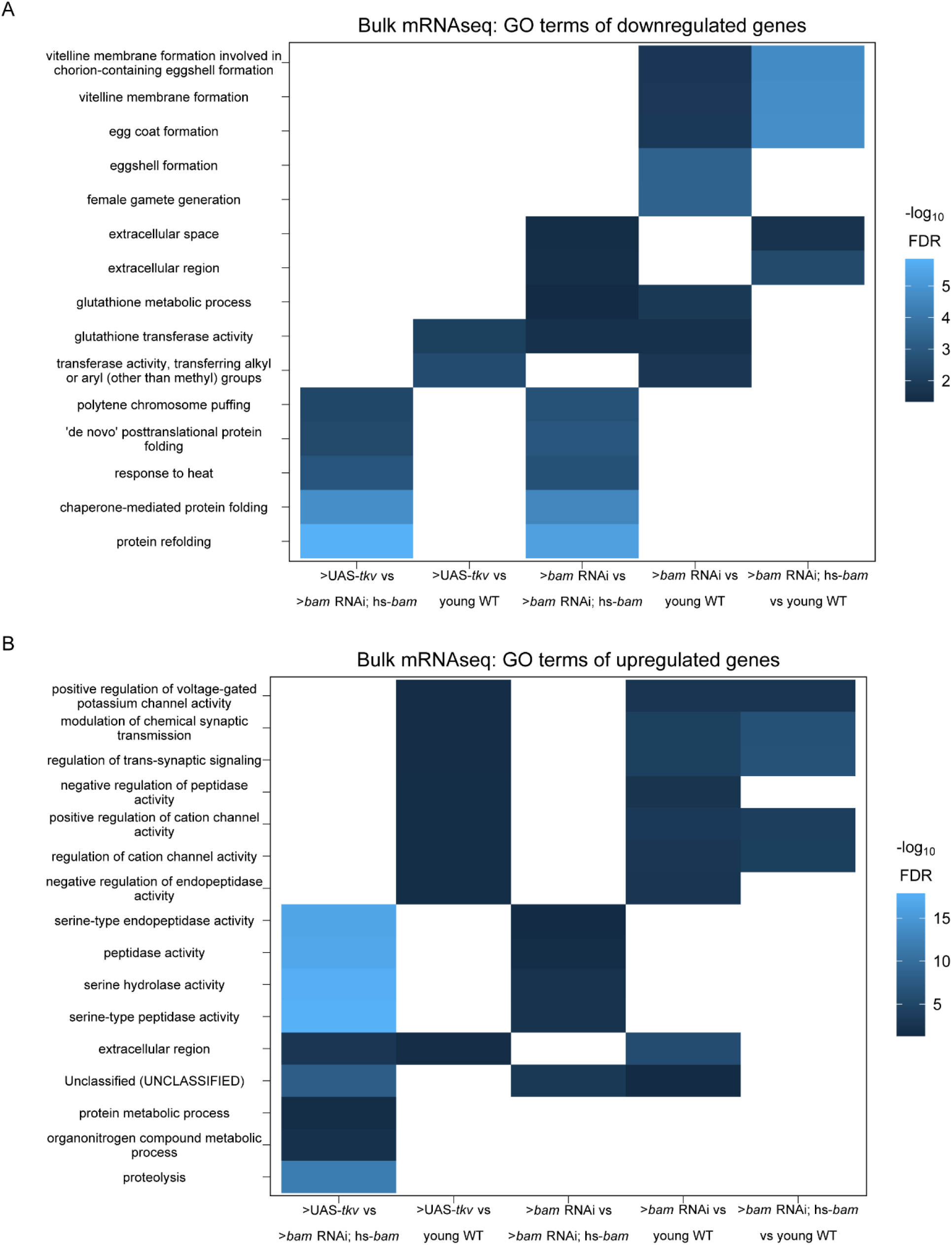
GO-terms enriched from differentially expressed genes between genetically enriched developmental milestones. (**A-B**) Heatmaps of top five significant GO-terms by fold enrichment resulting from each pairwise comparison of significantly (A) downregulated or (B) upregulated genes in the first genotype listed relative to the second genotype listed in the x-axis from bulk RNA-seq of each developmentally enriched stage. Comparisons that did not generate any significant GO-terms are omitted.

Next, to determine if our data could resolve large-scale expression changes that occur during oogenesis we examined the expression of genes in the GO-term meiotic cell cycle. Meiosis is initiated during the cyst stages of differentiation and therefore we would expect genes in the category, in general, to increase in expression in the >*bam* RNAi; hs-*bam* samples (Carpenter, 1979; Tanneti et al., 2011). We were surprised to find no significant change in the mean mRNA expression of genes in this GO-term in any of our enriched stages compared to enriched GSCs, though this does not preclude gene expression changes for individual genes (**Supplemental Figure 3A**). However, this is consistent with the observation that several factors that promote meiosis I are transcribed in the GSCs and the cells that follow (McCarthy et al., 2021). This suggests that, in general, a transition from a mitotic state to a meiotic state is not driven by large changes in mRNA levels of meiotic genes.

As we did not see overall changes to mRNA levels of genes in the GO-term meiotic cell cycle, we next examined the polysome-seq data of those genes to determine if changes in expression might occur at the level of translation. Polysome-seq uses polysome profiling to separate mRNAs that are associated with polysomes which form by mRNAs engagement with multiple ribosomes. To quantify the degree to which an mRNA is associated with polysome fractions, we compared the relative abundance of mRNAs from the polysome fractions to their relative expression using corresponding input lysates to calculate a metric referred to as translational efficiency (TE). Indeed, genes in the meiotic cell cycle GO-term had a significant increase in translation efficiency in CBs and a more dramatic increase in cysts despite no significant changes to the overall mRNA level of these genes (**Supplemental Figure 3A-B**). Based on scRNA-seq data, the expression of meiotic cell cycle genes increased slightly but significantly in the 4-CC cluster with a median increase in expression of 1.25 fold (**Supplemental Figure 3C**). This suggests that some genes in the meiotic cell cycle GO-term may be regulated at the mRNA level, but as a group this regulation is modest. This is likely because genes in this GO-term are robustly expressed even in GSCs as the median mRNA level of meiotic cell cycle genes in enriched GSCs is 36.1 TPM, which exceeds the 70^th^ expression percentile among all genes in enriched GSCs.

To validate this finding, we examined *orientation disrupter* (*ord)* because it is a well-characterized gene, is required for sister chromatid cohesion, and has previously been reported to peak in expression as meiosis begins in *Drosophila* (Bickel et al., 1997, 1996; Khetani and Bickel, 2007). Our Oo-site results suggested that *ord* mRNA was expressed before meiosis, both from bulk RNA-seq (**Figure 4A**) and scRNA-seq (**Supplemental Figure 3D**) consistent with reports that chromosome pairing initiates before meiotic entry (Christophorou et al., 2013; Joyce et al., 2013). However, polysome-seq data were consistent with the observation that Ord protein expression increases during the cyst stages due to translation (**Figure 4B**). This led us to predict that *ord* mRNA would be expressed before meiosis, and that Ord protein expression would increase during the cyst stages as previously observed, implying a change in the translation status of *ord* mRNA. To test this, we performed fluorescent *in situ* hybridization against GFP in a fly expressing Ord-GFP under the control of the *ord* promoter and 5’UTR. We visualized both the GFP protein and the mRNA and observed increased expression of Ord::GFP protein but consistent *ord::GFP* mRNA expression, indicating that Ord is controlled post-transcriptionally, likely at the level of translation based on our polysome-seq data (**Figure 4C-D’**). This finding also underscores the utility of Oo-site in exploring post-transcriptional gene expression changes.

**Figure 4:**
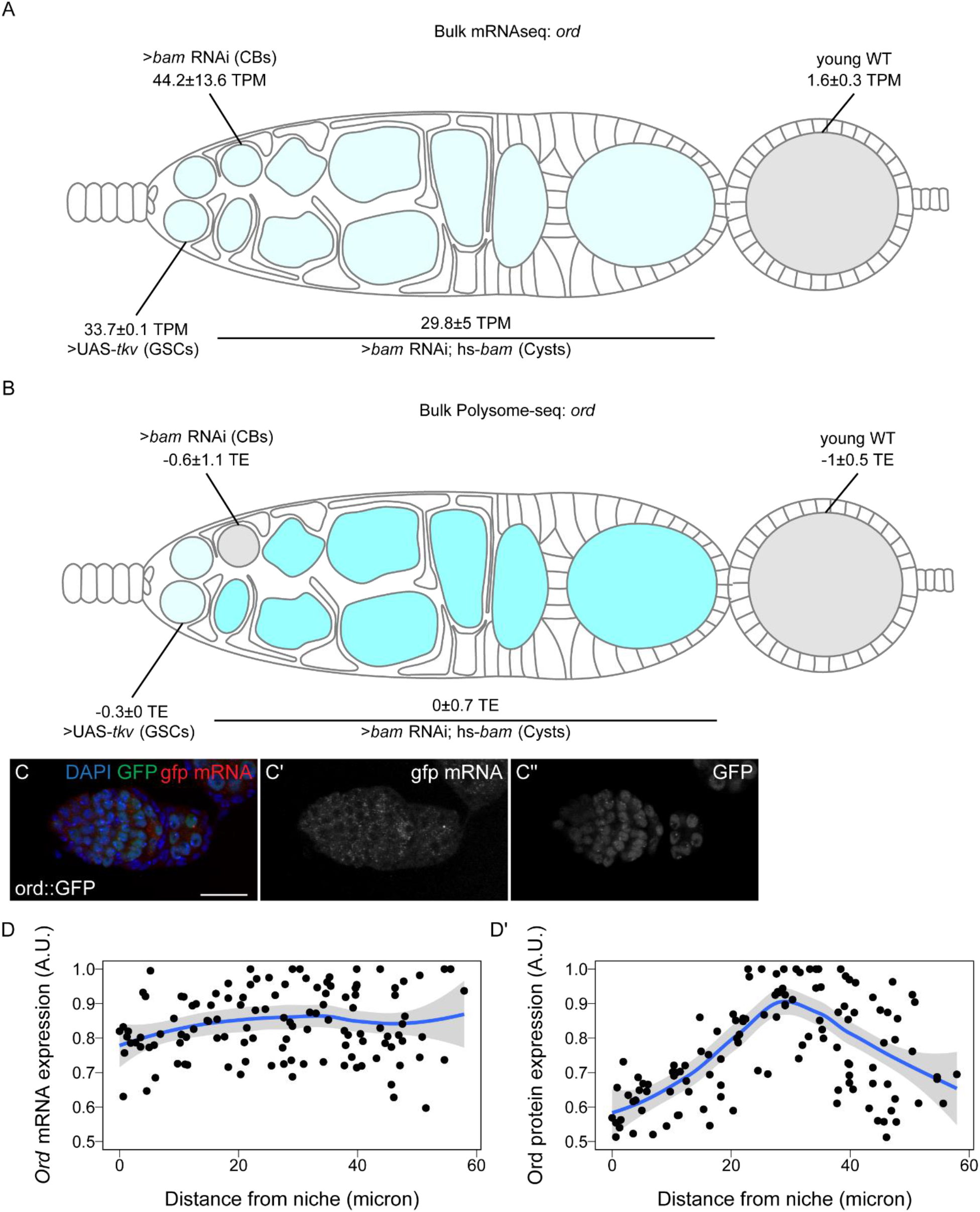
Ord expression is controlled post-transcriptionally. (**A-B**) Visualization of expression of *ord* over germline development from (A) bulk RNA-seq of developmentally enriched stages and (B) polysome-seq of developmentally enriched stages indicates that the mRNA level of *ord* is consistent from GSCs to cysts, until decreasing in early egg chambers, but the translation efficiency of *ord* increases during the cyst stages compared to other stages. Color indicates (A) relative expression or (B) TE and values indicate the (A) mean TPM±standard error or (B) the log_2_ mean TE±standard error (**C-C’’**) Confocal images of ovaries expressing Ord::GFP with *in situ* hybridization of *gfp* mRNA (red, middle greyscale) and stained for GFP protein (green, right greyscale) and DAPI (blue) demonstrate that the mRNA level of Ord::GFP is consistent throughout the germarium. (**D-D’**) Quantification of normalized mean intensity of stainings (C-C’’). X-axis represents the distance in microns from the niche, Y-axis represents mean intensity normalized to the maximum mean intensity per germarium of *ord* mRNA (D) or Ord protein (D). The line represents fit using a loess regression, shaded area represents the standard error of the fit. (n=8 germaria).

To further determine if meiosis is regulated post-transcriptionally, we examined the expression of genes in the GO-term “Double-strand break repair”, which is known to occur during meiosis 1 (Hughes et al., 2018; Page and Hawley, 2003). Double-stranded breaks are resolved before egg chamber formation (Hughes et al., 2018; Mehrotra and McKim, 2006; Page and Hawley, 2003). At the level of input mRNA, we found no significant changes in the expression of genes in this category compared to enriched GSCs (**Figure 5A**). From scRNA-seq data, the median expression of double-strand break repair genes significantly increases, but the median increase was only 1.05 fold in 4-CCs and 1.06 in 8-CCs compared to the GSC/CB/2CC group (**Figure 5B**). This suggests that double-strand break repair gene transcription begins in GSC stages and increases modestly during the cyst stages.

**Figure 5.**
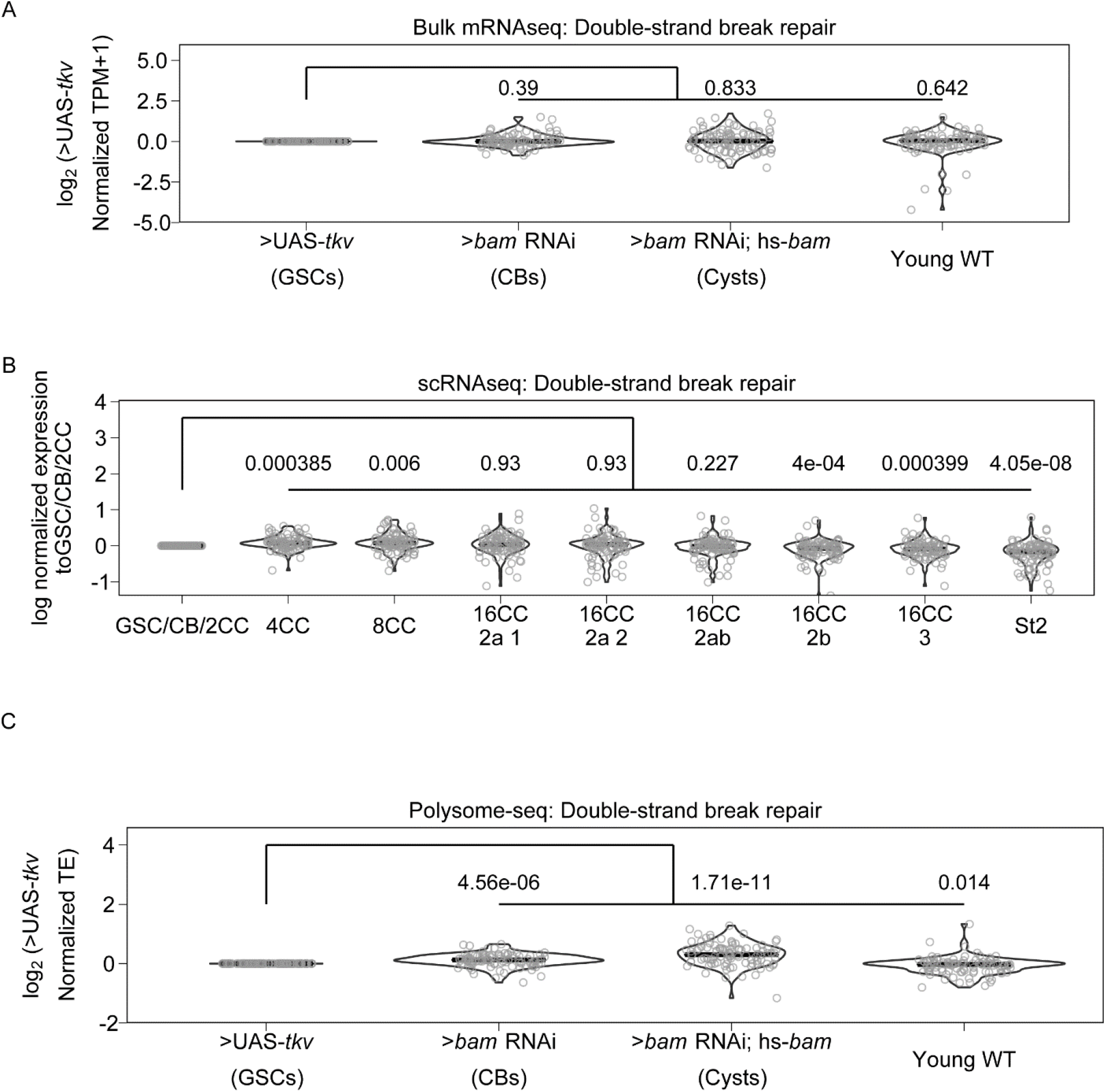
Genes involved in double-strand break repair may be controlled post-transcriptionally. (**A**) Violin plot of expression of genes in the GO category “Double-strand break repair” from bulk RNA-seq. No significant overall change in expression of these genes occurs comparing each genetically enriched developmental stage to GSCs. (**B**) Violin plot of expression of genes in the GO category “Double-strand break repair” from scRNA-seq. Overall expression of these genes increases in CBs, cysts, and young-WT ovaries compared to the GSC/CB/2CC cluster. Values above plots represent Holm-Bonnferroni adjusted p-values from a Welch’s t-test between the indicated genotypes (**C**) Violin plot of expression of genes in the GO category “Double-strand break repair” from polysome-seq. Overall expression of these genes increases in CBs, cysts, and young-WT ovaries compared to GSCs. Values above plots represent Holm-Bonnferroni adjusted p-values from a Welch’s t-test between the indicated genotypes.

In contrast, we found a significant increase in the median translational efficiency of double-strand break repair genes, with a 1.20 fold increase in the median translational efficiency in enriched CBs and a 1.56 fold increase in enriched cysts compared to enriched GSCs (**Figure 5C**). In young-WT the median fold change in translational efficiency decreased slightly but significantly compared to enriched GSCs at 0.95 fold. This is consistent with the observed progression of double-stranded break repair that occurs *in vivo*. This demonstrates that Oo-site can be used to derive insights into biological processes that may be changing during early oogenesis (Mehrotra and McKim, 2006; Page and Hawley, 2003). That key processes related to meiosis and differentiation are controlled post-transcriptionally is consistent with the importance of proteins that regulate translation such as Bam and Rbfox1 in differentiation and meiotic commitment during *Drosophila* oogenesis (Blatt et al., 2020; Carreira-Rosario et al., 2016; Flora et al., 2018; Kim-Ha et al., 1995; Li et al., 2009; Slaidina and Lehmann, 2014; Tastan et al., 2010).

## Discussion

We have developed an application that facilitates analysis of bulk RNA-seq, sc RNA-seq, and polysome-seq data of early *Drosophila* oogenesis that is accessible to non-bioinformaticians. We have demonstrated its utility in representing expression at the mRNA and translation level. Additionally, we have demonstrated that it can be used to visualize the expression of groups of genes over development to facilitate hypothesis development. As with all sequencing data, care should be taken to validate findings from Oo-site as sequencing can be influenced by a myriad of factors.

We have used Oo-site to discover that key meiosis regulators such as proteins of the synaptonemal complex and proteins of the double-strand break machinery are regulated at the level of translation. This adds to our understanding of the mechanisms regulating the mitotic to meiotic transition. In future work, identifying the factors mediating the widespread post-transcriptional regulation of crucial meiotic genes and mechanistically how it drives the mitotic to meiotic transition is of high importance.

High-throughput sequencing has enabled researchers to generate more data than ever before However, the development of analysis tools that are usable without bioinformatics training that enable users to make sense of these data to generate hypotheses and novel discoveries has lagged (Shachak et al., 2007). Oo-site allows for hypothesis generation and discovery using the powerful model system of *Drosophila* oogenesis. We believe Oo-site might also have utility as a teaching and demonstration tool to introduce students to the power of genomics in developmental biology. The open-source nature of this software facilitates future tool development, which will be crucial as more researchers delve into more data-intensive scRNA-seq, where visualization tools are limited and produce plots that may be difficult to interpret for those not versed in bioinformatics. Oo-site can be supplemented in the future to include additional data such as Cut and Run for various chromatin marks, nascent mRNA transcription using transient transcriptome sequencing or similar techniques, or protein levels from mass-spectroscopy to further extend its utility in hypothesis development.

## Acknowledgements

We thank the Drs. Ruth Lehmann and Maija Sladina for sharing scRNA-seq data with us before publication of the manuscript. We are grateful to all members of the Rangan laboratory for discussion and comments on the manuscript. We thank Noor Kotb for naming the dashboard Oo-site. We also thank Dr. Florence L. Marlow for critically reading and editing the manuscript. P.R. is funded by the National Institutes of Health NIGMS (RO1GM11177 and RO1GM135628).

## Materials and Methods

The following RNAi stocks were used in this study; *ord-GFP* (Bickel Lab), *Rps19b::GFP* (McCarthy et al., 2021), *UAS-Dcr2;nosGAL4* (Bloomington stock #25751), *bam* RNAi (Bloomington #58178), *hs-bam*/TM3 (Bloomington #24637),

### Sequencing data

Polysome-seq data were obtained from previous studies conducted by the Rangan lab. Data are available via the following GEO accession numbers:

>UAS-*tkv* GSE171349

>*bam* RNAi GSE171349, GSE166275

>*bam* RNAi; hs-*bam* GSE143728

Young-WT GSE119458

Single-cell sequencing data were obtained from Slaidina et al., GEO accession: GSE162192

### Code Availability

All code used in the preparation of this manuscript is available on GitHub at https://github.com/elliotmartin92/Developmental-Landscape/tree/master/Paper

The codebase underlying Oo-site is available on GitHub at https://github.com/elliotmartin92/Developmental-Landscape/tree/master/ShinyExpresionMap

### Antibodies

Mouse anti-1B1 1:20 (DSHB 1B1), rabbit anti-GFP 1:2000 (abcam, ab6556), rabbit anti-Vasa 1:4000 (Upadhyay et al., 2016), chicken anti-Vasa 1:4000 (Upadhyay et al., 2016)

### Polysome-seq

Flies ready for heat shock were placed at 37°C for 2 hours, moved to room temperature for 4 hours, and placed back into 37°C for 2 additional hours. Flies were then left overnight at room temperature and the same heat shocking procedure was repeated for a total of 2 days. Flies were then dissected in 1x PBS. Polysome-seq was performed as previously described (McCarthy et al., 2021).

### Polysome-seq data processing

Reads were mapped to the *Drosophila* genome (dm6.01) using STAR version 2.6.1c. Mapped reads were assigned to features also using STAR. Translation efficiency was calculated as in (Flora et al., 2018) using an R script which is available in the Oo-site Github repo. Briefly, TPMs (transcripts per million) values were calculated The log_2_ ratio of TPMs between the polysome fraction and total mRNA was calculated as such to prevent zero counts from overly influencing the data and to prevent divide by zero errors: 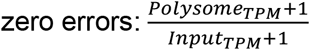. This ratio represents TE, TE of each replicate was averaged and standard error about the calculated average for each gene was calculated.

### Differential Expression

Differential expression analysis between all bulk RNA-seq samples in a pairwise manner was performed using DEseq2 (Love et al., 2014). Differential expression was considered as Foldchange > |4| fold, FDR < 0.05.

Differential expression analysis between all polysome-seq samples in a pairwise manner was performed using DEseq2 (Love et al., 2014) using the model ∼ type + genotype + genotype:type with LRT (reduced = ∼ type + genotype) to test for changes in polysome counts controlling for input counts. Differential expression was considered as (Foldchange > |2| fold, pvalue < 0.05)

Differentially expressed genes between all germline clusters from scRNA-seq was determined using the FindAllMarkers function from Seurat (Hao et al., 2021). Cutoff was logfc.threshold = 0.75.

Differentially expressed genes between all germarium soma clusters from scRNA-seq was determined using the FindAllMarkers function from Seurat (Hao et al., 2021). Cutoff was logfc.threshold = 0.75.

### GO term heatmaps

GO-term enrichment analysis was performed using Panther (release 20210224) using the default settings for an Overrepresentation Test of genes differentially expressed between Input samples. Top 5 GO-terms based on fold enrichment of each category were plotted using ggplot2 (Wickham, 2016).

### Fluorescent *in situ* hybridization

A modified *in situ* hybridization procedure for Drosophila ovaries was followed from Sarkar *et al*. (2021). Probes were designed and generated by LGC Biosearch Technologies using Stellaris® RNA FISH Probe Designer, with specificity to target base pairs of target mRNAs. Ovaries (3 pairs per sample) were dissected in RNase free 1X PBS and fixed in 1 mL of 5% formaldehyde for 10 minutes. The samples were then permeabilized in 1mL of Permeabilization Solution (PBST+1% Triton X-100) rotating in RT for 1 hour. Samples were then washed in wash buffer for 5 minutes (10% deionized formamide and 10% 20x SSC in RNase-free water). Ovaries were covered and incubated overnight with 1ul of the probe in hybridization solution (10% dextran sulfate, 1 mg/ml yeast tRNA, 2 mM RNaseOUT, 0.02 mg/ml BSA, 5x SSC, 10% deionized formamide, and RNase-free water) and primary antibody at 30°C. Samples were then washed 2 times in 1 mL wash buffer with 1ul of corresponding secondary antibody for 30 minutes each and mounted in Vectashield (VectaLabs).

### Quantification of Stainings

Stainings were quantified using the Fiji Measure tool. Images were aligned and cropped to place the stem cell niche at x=0. Individual cells were outlined within the germarium and Measure was used to calculate the Mean intensity of staining within the cell as well as the X coordinate of the centroid of the cell. Values were normalized to 1 by dividing Mean Intensity values by the maximum of the Mean Intensity per germarium. Data were plotted using ggplot2 and a fit line was added using ggplot2 geom_smooth with a “loess” function with default settings. The shaded area around the line represents standard error.

---

**Supplemental Figure 1.**
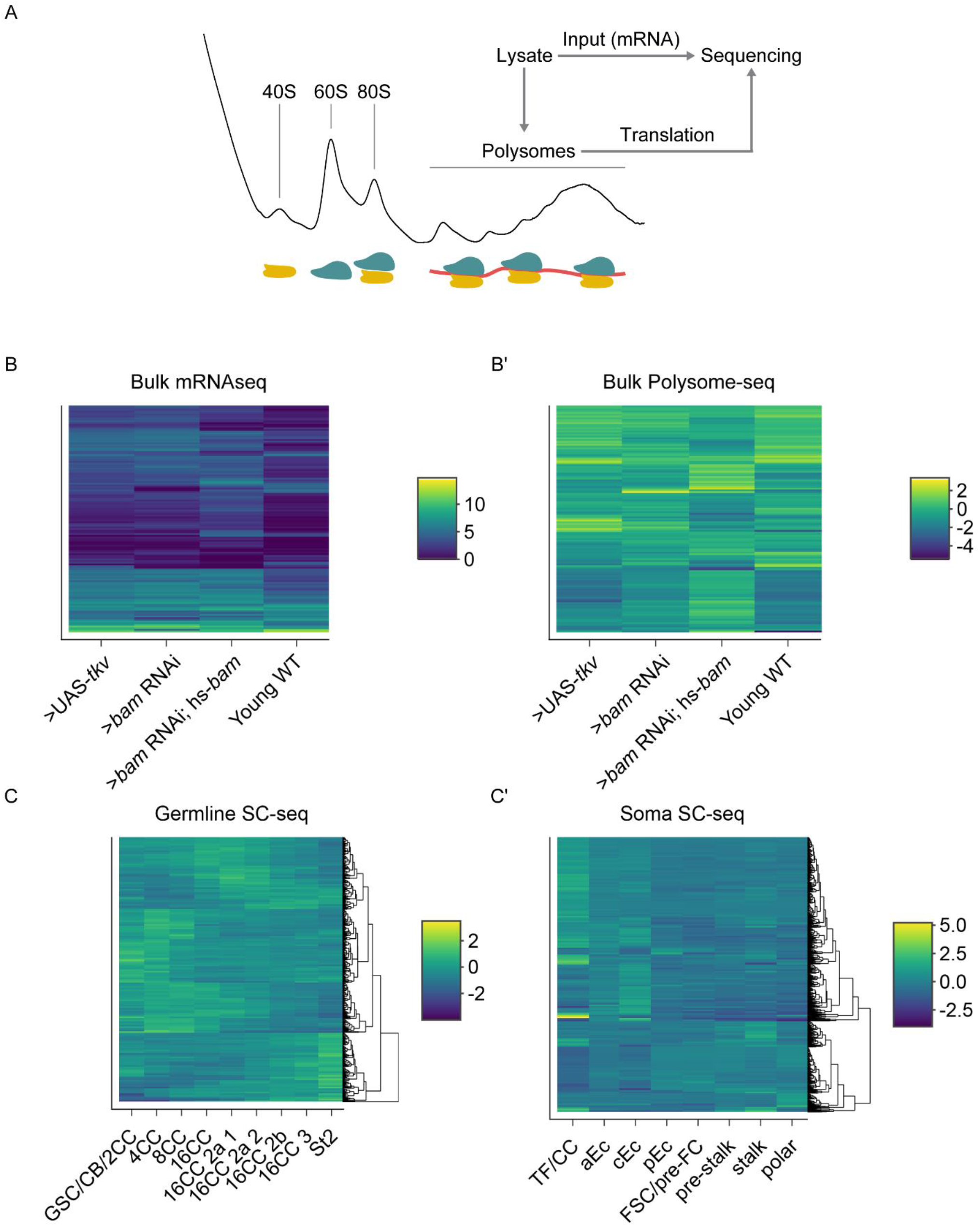
Sequencing strategy and clustered heatmaps of differential expression, related to Figure 1. (**A**) Schematic of strategy used to obtain input mRNA samples and matched polysome-seq libraries of ovaries genetically enriched for developmental milestones. (**B-B’**) Clustered heatmaps of (B) bulk RNA-seq and (B’) log_2_(TE) from bulk polysome-seq of the developmental milestones indicated on the X-axis. Each row in the heatmap indicates a gene that is differentially expressed in at least one of the milestones compared to all others in a pairwise fashion. Color scale denotes average relative expression. (**C**) scRNA-seq of early germline cells and (**C’**) scRNA-seq of somatic cells in the germarium. X-axis denotes cell-type and each row in the heatmap indicates a gene that is differentially expressed in at least one of the cell-types compared to all others in a pairwise fashion.

**Supplemental Figure 2.**
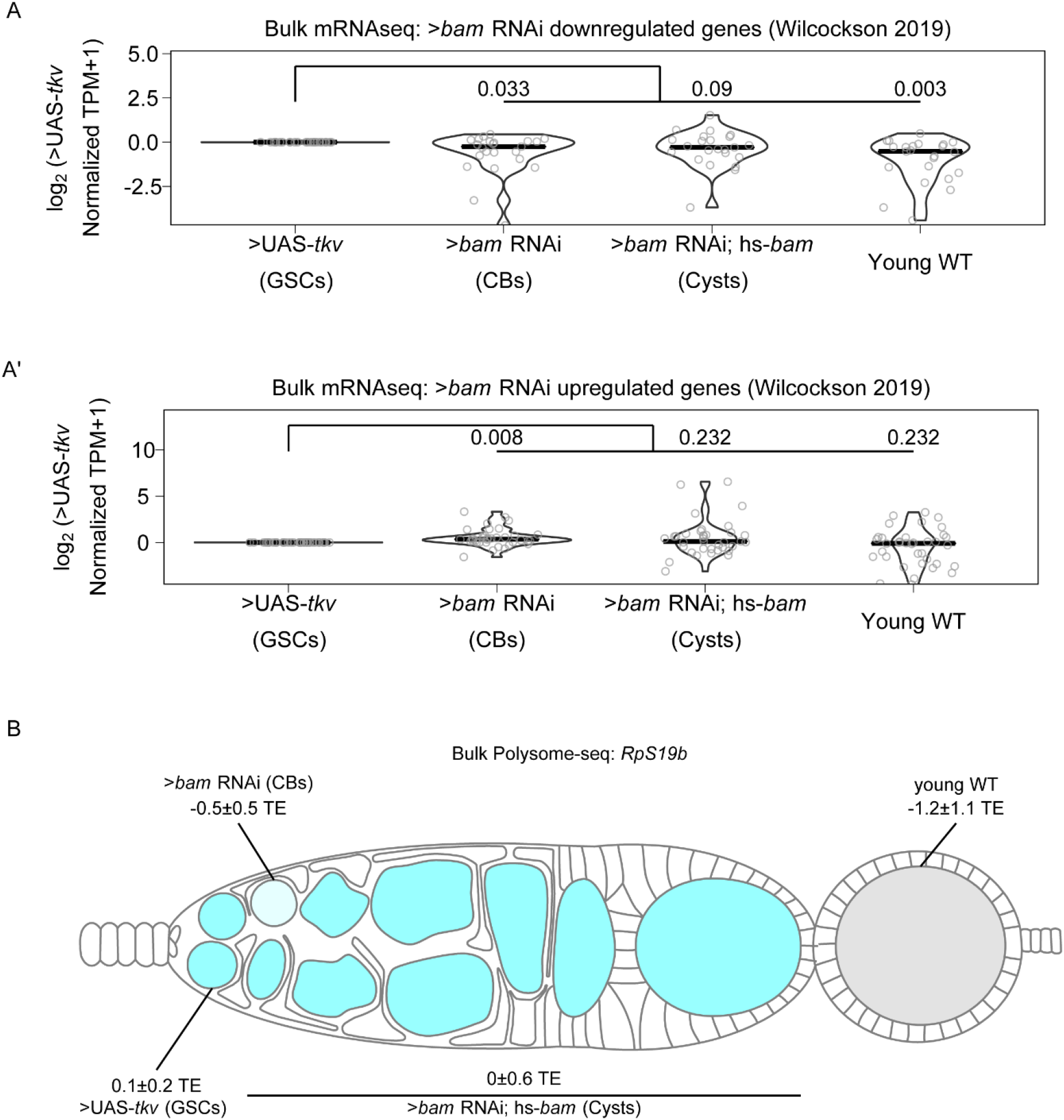
Bulk RNA-seq recapitulates previously observed expression patterns of gene expression, related to Figure 2. (**A-A’**) Violin plots of expression from bulk RNA-seq of genes 2-fold or more (A) down or (A’) upregulated in bam RNAi germline cells compared to UAS-TKV overexpressing germline cell with a p-value < 0.01 over germline development from Wilcockson *et al*. demonstrate that bulk RNA-seq identifies similar trends in gene expression compared to the FACS based method employed by Wilcockson *et al*. Values above plots represent Holm-Bonnferroni adjusted p-values from a Welch’s t-test between the indicated genotypes. (**B**) Visualization of expression of *RpS19b* over germline development from polysome-seq data. Color indicates TE and values indicate the log_2_mean TE±standard error *RpS19b* TE is relatively consistent during early oogenesis and decreases in the egg chambers.

**Supplemental Figure 3.**
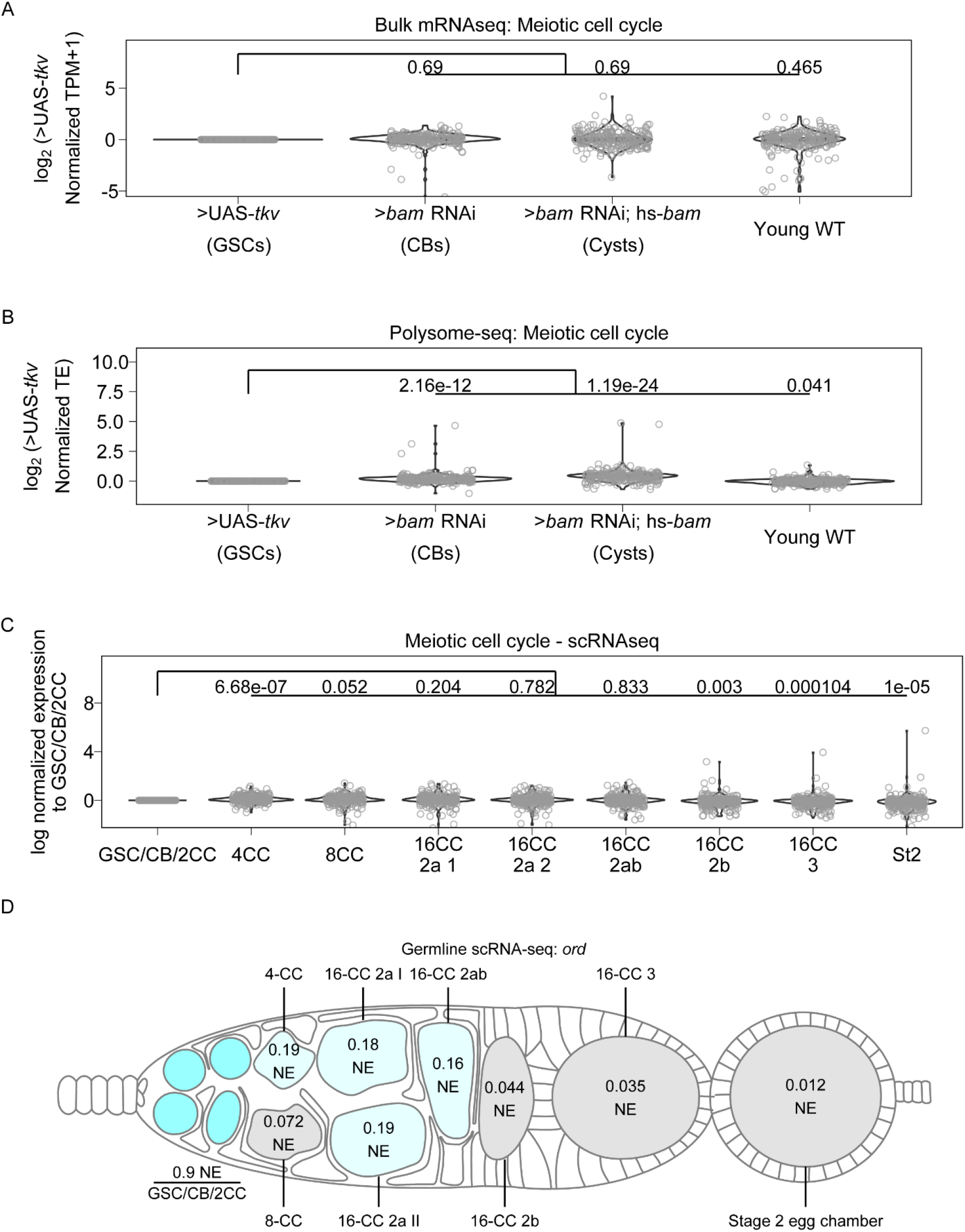
Genes involved in meiotic cell cycle, including *ord*, may be controlled post-transcriptionally, related to Figure 4. (**A**) Violin plots of gene expression from RNA-seq of genes in the GO-term category meiotic cell cycle. No significant overall change occurs to expression of these genes at any of the developmental milestones compared to GSCs. Values above plots represent Holm-Bonnferroni adjusted p-values from a Welch’s t-test between the indicated genotypes. (**B**) Violin plots of TE from polysome-seq of genes in the GO-term category meiotic cell cycle. Overall TE increases in CBs and cysts significantly compared to GSCs indicating that meiotic entry may be partially controlled post-transcriptionally. Values above plots represent Holm-Bonnferroni adjusted p-values from a Welch’s t-test between the indicated genotypes. (**C**) Violin plot of expression of genes in the GO category “meiotic cell cycle” from scRNA-seq. Overall expression of these genes increases in CBs, cysts, and young-WT ovaries compared to the GSC/CB/2CC cluster. Values above plots represent Holm-Bonnferroni adjusted p-values from a Welch’s t-test between the indicated genotypes. (**D**) scRNA-seq data indicate that the mRNA level of *ord* is highest in the GSC/CB/2CC cluster, but remains relatively consistent in its expression starting in the 4-CC through 16-CC 2ab clusters and is dramatically decreased in early egg chambers. Color and values indicate the normalized expression of *ord* in each given stage.

